# Integrating image-based phenotyping and GWAS to map tolerance to Spittlebug nymphs in interspecific *Urochloa* grasses

**DOI:** 10.1101/2025.11.03.686394

**Authors:** Paula Espitia-Buitrago, Claudia Perea, Juan C. Mejia-Medina, Luis M. Hernández, Valheria Castiblanco, Camilla Ryan, José J. De Vega, Rosa N. Jauregui

## Abstract

*Urochloa* grasses are among the most widely used forage grasses across the tropics. Spittlebugs (*Hemiptera*: Cercopidae) are major pests of tropical *Urochloa* (syn. *Brachiaria*) pastures, severely reducing forage productivity and quality. Understanding the genetic basis of host-plant resistance is essential for developing durable resistant cultivars. Here, we combined high-throughput image-based phenotyping and genome-wide association studies (GWAS) to dissect the genetic architecture of tolerance to *Aeneolamia varia* nymphs in 339 interspecific F₁ hybrids derived from crosses between resistant sexual and susceptible apomictic *Urochloa* parents. Digital image analysis using both unsupervised (DQU) and supervised (DTR) quantification pipelines enabled precise estimation of plant damage, yielding moderate to high broad-sense heritability estimates (H² = 0.49-0.66). In contrast, insect survival (NTS) exhibited low to moderate correlations with all damage traits and lower heritability estimates (H² = 0.42). Using 57,051 high-quality SNPs aligned to the genome of the hybrid cultivar Basilisk, GWAS models identified 18 quantitative trait loci (QTL) for plant damage traits, but none for insect survival (antibiosis). Six robust QTL on chromosomes 1, 6, 7, 27, 29, and 36 were consistently detected across models and phenotyping methods, explaining up to 21.5% of phenotypic variance. Candidate gene analysis revealed proteins involved in hormone signalling, oxidative stress response, and cell wall modification, suggesting multifaceted tolerance mechanisms. These results provide a foundational set of molecular markers associated with spittlebug tolerance in *Urochloa,* useful for marker-assisted and genomic selection in our forage breeding programme.

## Introduction

*Urochloa* grasses are among the most widely used forage grasses across the tropics. Spittlebugs (*Hemiptera*: Cercopidae) are a major pest of *Urochloa* P. Beauv. grasses (syn. *Brachiaria*) in tropical America, causing significant plant damage, particularly during the rainy season. Their infestations not only reduce forage production but also negatively impact nutritional quality, including crude protein content and *in vitro* digestible dry matter, resulting in reductions in livestock weight gain up to 74% per hectare (Congio et al., 2020). Given that pasture productivity directly impacts animal productivity in the dairy and beef production chains, host plant resistance is the cornerstone of integrated pest management in *Urochloa* grass pasture systems.

Host plant resistance is a particularly advantageous strategy against pests because it maintains pest populations below the economic injury threshold, it is fully compatible with other control methods, and it is inherently delivered through the seed, reducing long-term management costs (Cardona & Sotelo, 2005). A resistant plant is defined as one that, under pest pressure, exhibits reduced damage as a result of its genetic makeup, with this resistance determined by various underlying mechanisms (i.e., traits that comprise the expression of resistance) and grouped into the resistance categories of antibiosis, antixenosis, and tolerance (Souza, 2025; Stout et al., 2024). Nonetheless, breeding for insect resistance remains difficult due to the high genetic polymorphism of the phytopathogenic agents, and being strongly shaped by environmental conditions and regulated by host–pathogen interactions (Priyadarshan, 2019). Thus, identifying the resistance categories involved in genotypes’ responses to herbivory is crucial to diversifying the basis of resistance through diverse loci and, ultimately, building long-term resistance in breeding populations (Pamidi et al., 2025; Peterson et al., 2017).

Breeding programs for *Urochloa* aim to incorporate resistance traits present in diverse genotypes to develop improved cultivars. Notably, spittlebug resistance in *Urochloa* is both interspecific, where host genotype’s response varies among spittlebug species, and intraspecific, where resistance depends on the insect’s developmental stage (Cardona et al., 2010). To the best of our knowledge, the molecular mechanisms of resistance to spittlebugs have not been clearly elucidated in *Urochloa*. However, several factors have been proposed in past studies, including the production of volatile organic compounds (Silva et al., 2017, 2019), improved tolerance response with fertilisation (Alvarenga et al., 2019), and activation of defence-related biochemical responses such as lipoxygenases, proteases, jasmonic acid and abscisic acid (Barros et al., 2021).

Despite these findings, translating such resistance traits into practical outcomes through molecular breeding approaches in *Urochloa* grasses remains a significant challenge. The diverse reproductive modes, both apomictic and sexual, within intra- and interspecific crosses, a broad range of ploidy levels, and the outcrossing nature of *Urochloa* grasses, combined with limited understanding of chromosomal behaviour during meiosis, unclear modes of inheritance, and high heterozygosity, significantly hinder crossing, evaluation and selection activities (Ferreira et al., 2021). Despite these challenges, F₁ families have been utilised to identify quantitative trait loci (QTL) linked to key agronomic traits such as apomixis (Worthington et al., 2016), aluminum tolerance (Worthington et al., 2021), forage yield (Vidotti et al., 2019), and spittlebug antibiosis (Ferreira et al., 2019). The implementation of molecular markers in crossing, evaluation, and selection activities remains limited, with notable success primarily in marker-assisted selection of apomictic plants using markers p779/p780 (da Costa Lima Moraes et al., 2023; Nitthaisong et al., 2019; Worthington et al., 2016).

Previous QTL mapping studies in *Urochloa* have relied on the diploid *U. ruziziensis* reference genome, which may not fully capture the genomic complexity of the polyploid interspecific hybrids commonly used in breeding programs. Recently, the allotetraploid reference genome of *cv.* Basilisk (CIAT 606) from the interspecific *Urochloa* complex has been released and found to be a hybrid of *U. decumbens* and *U. brizantha* (Ryan et al., 2025). It provides a more appropriate genomic framework for QTL analysis in polyploid breeding materials. Given the complexity of spittlebug resistance and the limited understanding of its genetic architecture, implementing this improved genomic resource for QTL mapping can provide enhanced resolution for trait dissection and facilitate the identification of candidate genes underlying resistance mechanisms. Such insights are crucial for developing functional markers that could be implemented in breeding programs and for understanding the biological basis of host-plant resistance to spittlebugs.

The objective of this study was to identify genomic regions associated with host-plant resistance to *Aeneolamia varia* (Fabricius, 1787) (Hemiptera: Cercopidae) nymphs in interspecific *Urochloa* hybrids and explore candidate genes that may contribute to understanding the resistance mechanisms.

## Methods

### Plant material and phenotypic data

Four tetraploid interspecific sexual hybrids (*U. ruziziensis* x *U. brizantha* x *U. decumbens*) resistant to *A. varia* were crossed with the common susceptible apomictic tester *Urochloa decumbens* cv. Basilisk (CIAT 606; genome references), comprising four F₁ biparental families with a total of 339 genotypes (Table 1).

**Table 1.** Biparental F_1_ families of *Urochloa* interspecific hybrids.

| Cross | Number of individuals of the progeny |
| --- | --- |
| Sx14_808 x CIAT 606 | 70 |
| Sx14_793 x CIAT 606 | 120 |
| Sx14_123 x CIAT 606 | 120 |
| Sx14_1042 x CIAT 606 | 29 |

The F₁ progeny was evaluated for resistance to nymphs of *Aeneolamia varia* (Hemiptera: Cercopidae) in no-choice tests under controlled greenhouse conditions using two commercial cultivars as resistant controls (*U.* interspecific cv. Mulato II, and *U. brizantha* cv. Marandú) at the Center for Tropical Agriculture (CIAT), Palmira, Colombia (3°30’12.34“N, 76°21’23.81“W). The no-choice tests were conducted in seven independent trials, T1 to T7, each following a Federer augmented-block experimental design with one infested and one uninfested replicate for the 339 hybrids and the five parentals. The controls were included in each incomplete block. Following Parsa et al. (2011), each single-stemmed experimental unit with exposed roots was infested with six *A. varia* eggs close to the hatching date. Treatment considered the factor genotype and the factor infestation.

Response variables scored included insect survivorship (%) (NTS) (Cardona et al., 2010) and plant damage (%). Both were evaluated at 35 days after infestation (DAI), which aligns with the nymphal stage duration of this spittlebug species (Fig. 1). NTS was evaluated in five trials, excluding T1 and T4, and was expressed as the total of living insects in nymphal and adult instars stages at 35 DAI:

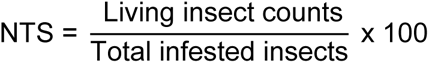

**Figure 1.**
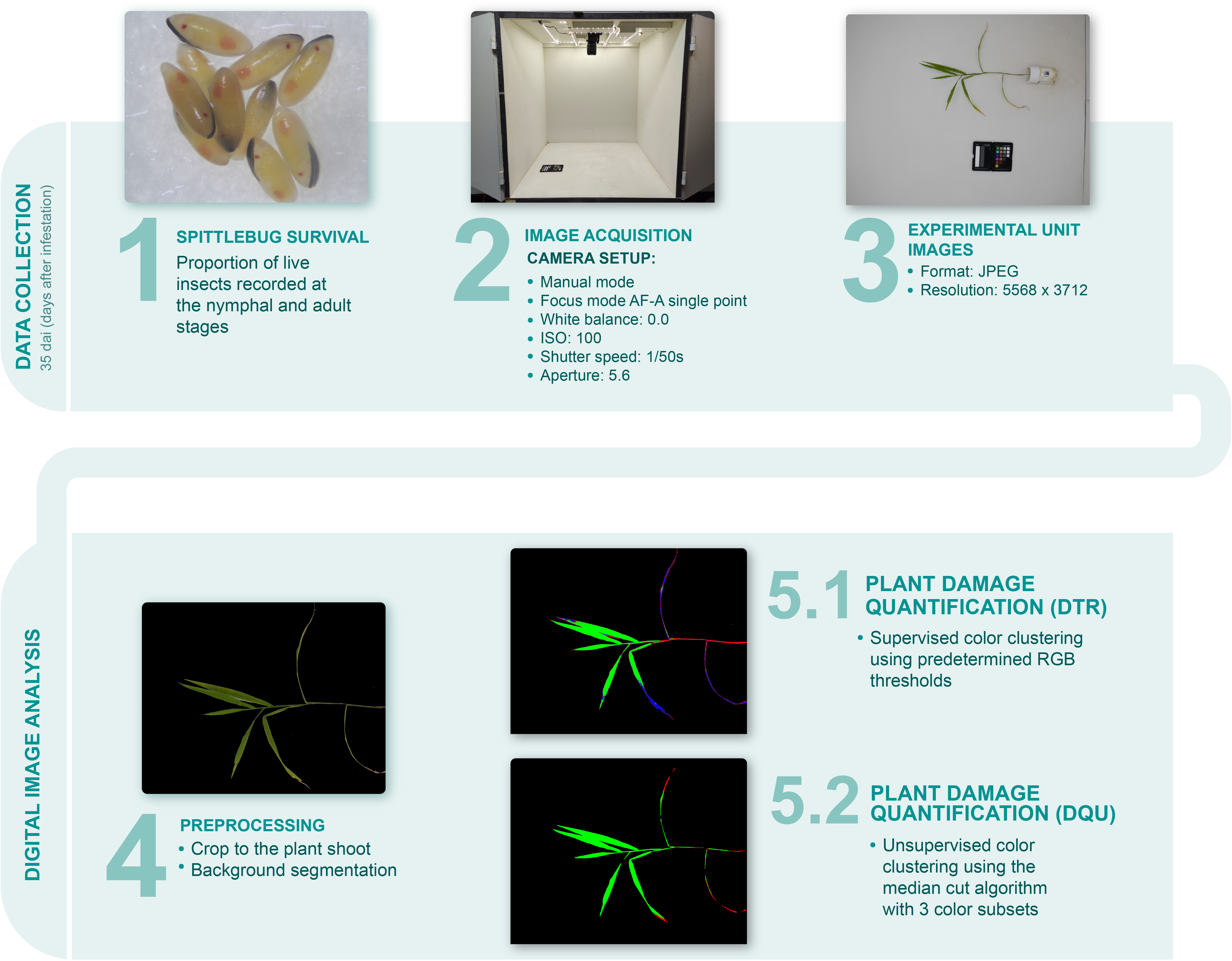
Response variables collected in no-choice trials to *Aeneolamia varia* nymphal infestation in interspecific *Urochloa* hybrids, and digital image analysis workflow to quantify plant damage.

Plant damage percentage was quantified across all trials from digital images acquired in a lightbox under controlled lighting conditions. Images were captured using a Nikon D7500 reflex camera with fixed photographic parameters: AF-A single point focus, white balance 0.0, ISO 100, shutter speed 1/50 s, and aperture f/5.6. Quantification employed two separate methodologies. The first, Color Clustering Image Analysis (DQU), utilized an unsupervised color clustering algorithm within ImageJ (Ferreira & Rasband, 2012; Hernández et al., 2022). The second, RGB Pixel Counting (DTR), involved using the jpeg package in R (Espitia-Buitrago et al., 2025; Urbanek, 2011) to quantify green, chlorotic, and necrotic tissue based on predetermined Red, Green, and Blue (RGB) values. Tissue classification followed these specific criteria: Green tissue was defined by R < (0.9 × G) AND B < (0.9 × G) AND (2 × G) > (20/255) × (B+R) (Patrignani & Ochsner, 2015); Yellow (chlorotic) tissue was classified when (90/255) < R < (241/255) AND (97/255) < G < (216/255) AND (18/255) < B < (165/255); and Necrotic tissue was identified when (27/255) < R < (225/255) AND (56/255) < G < (213/255) AND (0/255) < B < (53/255). Finally, plant damage was quantified using the equation:

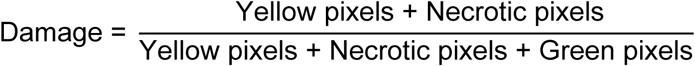

### Phenotypic Statistical analysis

Phenotypic data from the infested treatment were analysed using linear mixed models (LMM) implemented in ASReml-R version 4.3.3 (Butler et al., 2018). Plant damage traits, including total plant damage, were quantified with the unsupervised algorithm (DQU), total plant damage quantified with the supervised algorithm (DTR), yellow tissue and necrotic tissue quantified with the supervised algorithm (DTR). All metrics were evaluated across all trials. Insect survival (%) (NTS) was analyzed across five environments, excluding T1 and T4 where this variable was not recorded due to logistic constraints.

To ensure the data quality, each trail was assessed individually using an LMM with genotype fitted as a random effect to estimate Cullis broad-sense heritability based on the average pairwise prediction error variance between genotypes (Covarrubias-Pazaran, 2020). Trials with convergence issues or heritability estimates below 0.2 were excluded from further analysis. The final model incorporated trial (*i.e.* environment) as a fixed effect and was structured as:

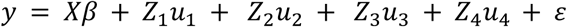

Where *X* is the design matrix and β the vector of trial fixed effects; *Z*_*i*_ are design matrices for *u*_*i*_ vectors: *Z*_1_corresponds to genotypic effects *u*_1_, assumed ∼ N(0, *A*σ^2^) using a pedigree-derived relationship matrix *A* (build via *A*-inverse using parental records of the hybrids); *Z*_2_ corresponds to genotype by trial random effects *u*_2_; *Z*_3_ corresponds to block random effects *u*_3_; *Z*_4_ corresponds to row x column spatial effects *u*_4_ . Residual errors (*ɛ*) were modeled as heterogeneous across trials using dsum(∼units | Trial).

To obtain Best Linear Unbiased Predictors (BLUPs) of genotypic performance across trials, genotype was modeled as a random effect. In contrast, Best Linear Unbiased Estimators (BLUEs) were calculated by treating genotype as a fixed effect. Cullis broad-sense heritability for each trait was estimated from the BLUP model variance components. Additionally, genotype-specific means were included by the arithmetic averaging of replicated measurements across trials, providing a preliminary measure of phenotypic performance. All estimates, including means, BLUEs, BLUPs, and variance components, were exported for downstream genome-wide association analyses (GWAS). Trait correlations and their significance were assessed using Pearson correlation coefficients calculated from BLUEs, as these represent genotypic values without shrinkage

### Genotypic data and bioinformatic workflow

Leaf samples of the 344 individuals were lyophilized for DNA extraction at Eurofins Genomics. Subsequently, these samples were processed by Floragenex Inc. for single-end RAD-seq using 118 bp reads with the PstI restriction enzyme. To ensure adequate coverage each F₁ individual was sequenced twice, while parental samples were sequenced eight times to achieve higher depth for accurate genotype calling. Raw reads were demultiplexed and quality-trimmed before mapping to the tetraploid *Urochloa decumbens* cv. Basilisk reference genome (GCA_964030465.3) (Ryan et al., 2025) using Bowtie2 algorithm version 2.1 (Langmead & Salzberg, 2012). Genotype calling was performed using NGSEP V.3 (Perea et al., 2016) with ploidy set to 2 (diploid mode), as previous studies suggest that mean read depths of at least 61 (Meireles, et al., 2019) are required for accurate tetraploid allele dosage estimation in *Urochloa* species, which is higher than the dataset coverage. The resulting VCF file was phased and imputed using Beagle (Browning et al., 2018, 2021) (version 5.5), followed by quality filtering to retain only biallelic SNPs with Minor Allele Frequency (MAF) ≥ 0.01 and Call Rate ≥ 40%. This filtering pipeline yielded 57,051 high-quality biallelic SNP markers for downstream analyses.

### Genome-Wide Association Study (GWAS) and Gene Annotation

GWAS analysis was conducted in GAPIT version 3 (Wang & Zhang, 2021), implementing two complementary algorithms: the Fixed and random model Circulating Probability Unification (FarmCPU) (Liu et al., 2016) and the Bayesian-information and Linkage-disequilibrium Iteratively Nested Keyway (BLINK) models (Huang et al., 2019). To account for population structure and relatedness, covariates included the first three principal components, and a kinship matrix calculated using the VanRaden method (VanRaden, 2008). Phenotypic data consisted of estimates per genotype derived from the LMM analyses for plant damage using both the DQU and DTR methodologies, as well as insect survival (%) (NTS). Genotypic data consisted of a quality-filtered VCF containing 57,051 biallelic SNPs.

Significant marker–trait associations (MTAs) were identified using a −log₁₀(P-value) threshold of 6 and validated through False Discovery Rate (FDR) correction using the Benjamini–Hochberg procedure, as implemented in GAPIT (Wang & Zhang, 2021). Quantitative trait loci (QTL) regions were then defined by applying three window sizes (±2 kb, ±5 kb, and ±10 kb) around each significant MTA, based on the estimated linkage disequilibrium (LD) decay distance (r² = 0.2) for each biparental populations. LD decay analysis was performed using PopLDdecay v1.4.4 (Zhang et al., 2019) with a minor allele frequency threshold of 0.01. PairwiseLD values (r²) were calculated between all SNP pairs and averaged across fixed genomic distance bins of 1 kb to generate decay curves per chromosome. The LD decay distance was estimated as the point where the fitted decay curve intersected r² = 0.2, for each F1 family and for the general population. Candidate genes located within the defined windows were extracted from the *U. decumbens* cv. Basilisk reference genome (GCA_964030465.3) (Ryan et al., 2025) and functionally annotated using BLASTp searches against the UniProt database (Bateman et al., 2025). Searches were conducted with default parameters, except for an E-value threshold of 0.01. Results were filtered to retain only hits with ≥85% identity, optimal alignment length coverage ≥70%, and high-confidence functional annotations (UniProt Protein Existence evidence levels 1 or 2: “Evidence at protein level” or “Evidence at transcript level”).

### Use of Large Language Models (LLMs)

Portions of the manuscript text and R code for data visualization were refined with assistance from ChatGPT (OpenAI GPT-4, version released July 2023, accessed via https://chat.openai.com/). The model was queried using natural language prompts to improve the clarity of scientific writing and to polish R code used for generating graphs and figures. All model outputs were critically reviewed, edited, and validated by the authors to ensure scientific accuracy, proper formatting, and originality.

## Results

### Phenotypic analysis

A high degree of phenotypic variation in plant damage and insect survival was observed among the *Urochloa* interspecific population in response to *Aeneolamia varia* nymph infestation. Infested plants showed higher levels of damage compared to non-infested plants across all the image-derived variables, regardless of the quantification method used, either DQU (unsupervised clustering with color quantization) or DTR (supervised clustering with RGB thresholds) (Fig. 2; Suppl. Fig. 1). Insect survival (NTS), calculated as the proportion of live insects 35 days after infestation, showed discrete distribution due to its derivation from fixed insect counts (Fig. 2).

**Figure 2.**
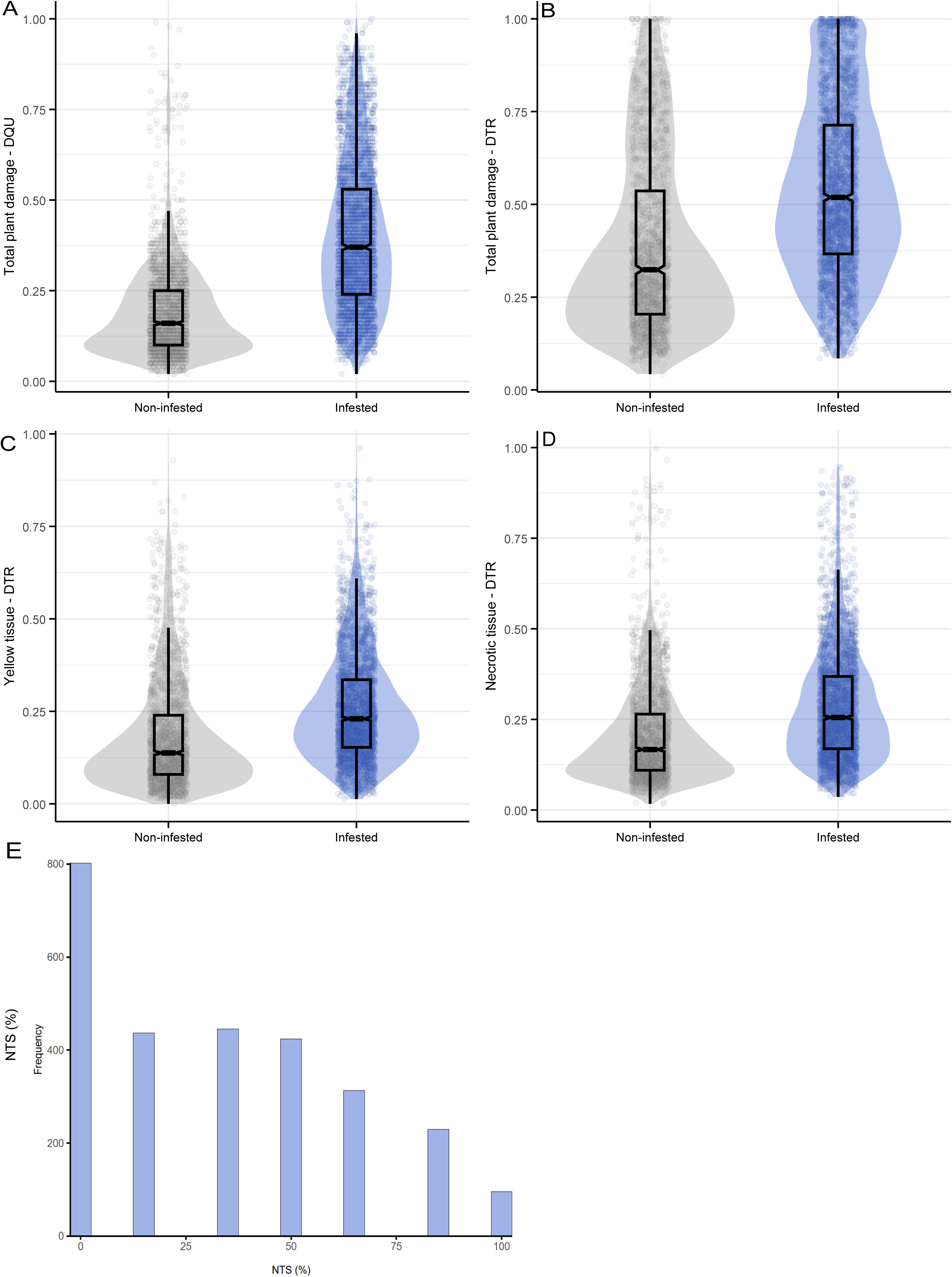
Violin plots displaying the distribution of raw plant damage variables and histogram displaying insect survival across the seven trials.

Trial-level analyses of broad-sense heritability revealed that trial T2 consistently showed low estimates (H² < 0.2) across all evaluated traits and was therefore excluded from final statistical analyses to ensure model stability and data quality (Suppl. Table 1). Notably, damage estimates were consistently higher with the DTR method compared to DQU; however, genotype classification remained consistent between both image analysis techniques (Fig. 3). A strong and highly significant correlation was observed between total plant damage estimates derived from DQU and DTR (r = 0.95), as well as with necrotic tissue (r = 0.81). Correlations between total damage and yellow tissue were slightly lower, ranging from r = 0.64 for the DQU method to r = 0.73 for DTR. In contrast, insect survival NTS exhibited low to moderate correlations with all damage traits (Fig. 4).

**Figure 3.**
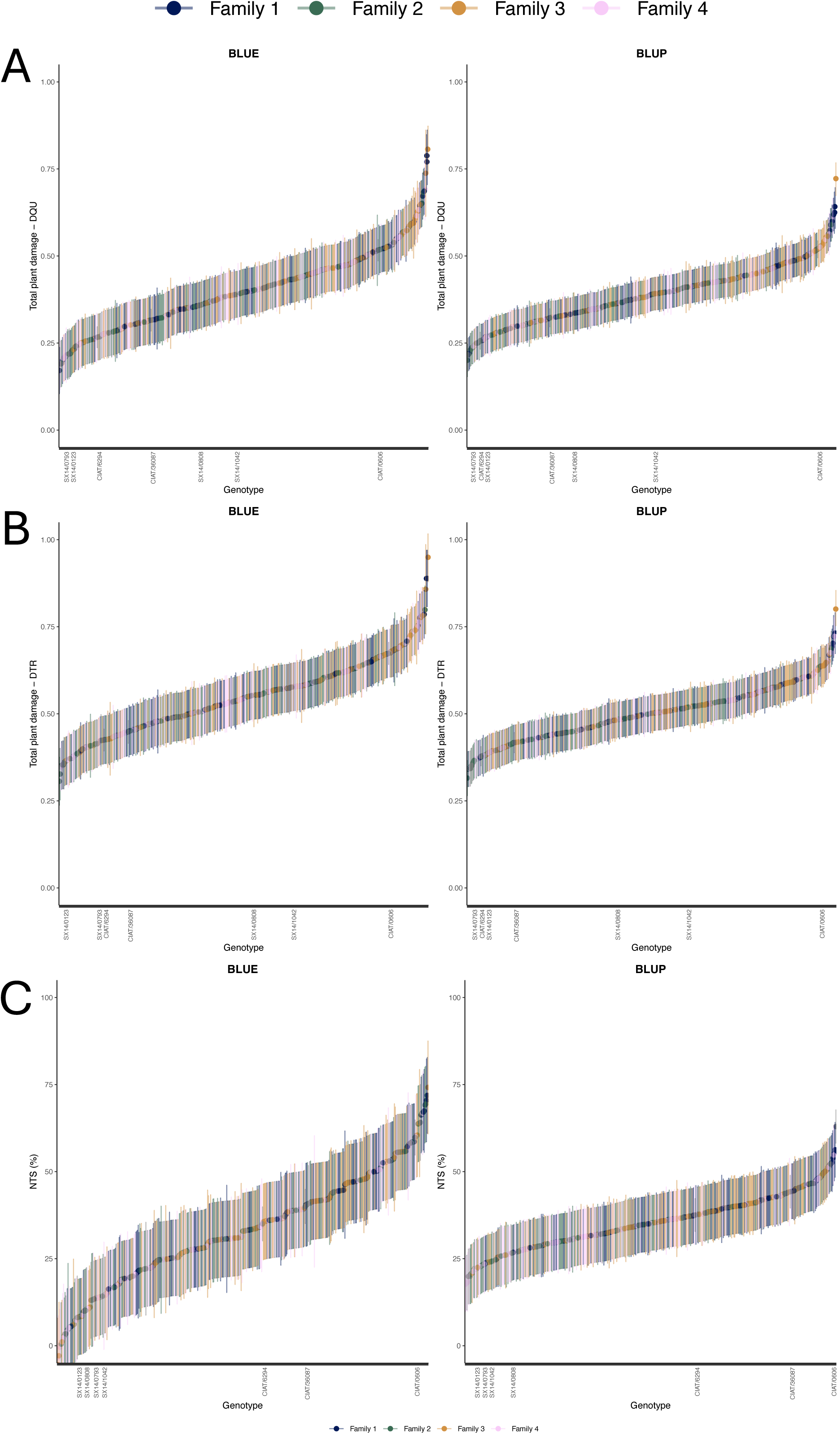
BLUEs and BLUPs of Total plant damage calculated with A) Unsupervised median cut algorithm for color quantization using three color subgroups (DQU); B) Supervised RGB threshold setting (DTR); C) Insect survival (NTS %).

**Figure 4.**
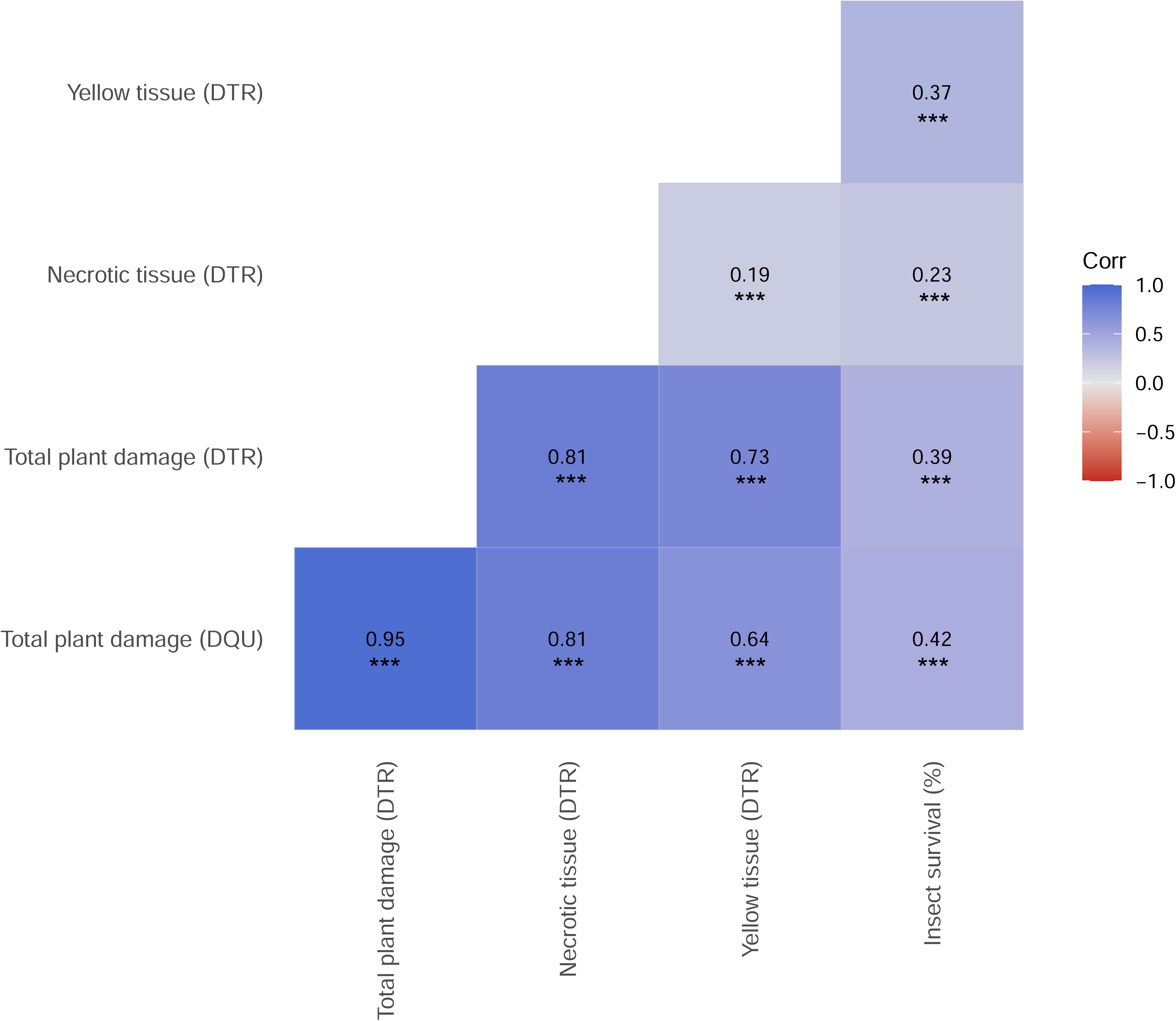
Pearson correlation matrix among BLUEs of plant damage traits and insect survival (%). Significance levels indicated by asterisks (*** = p-value < 0.001).

Broad-sense heritability estimates of plant damage traits ranged from moderate to high, with values from 0.49 for necrotic area (DTR) to 0.66 for total plant damage (DQU) (Table 2). NTS showed the lowest heritability estimate (0.42), reflecting the effect of its discrete, bounded nature and the high frequency of zero survival observed at 35 days after infestation (DAI).

**Table 2.**
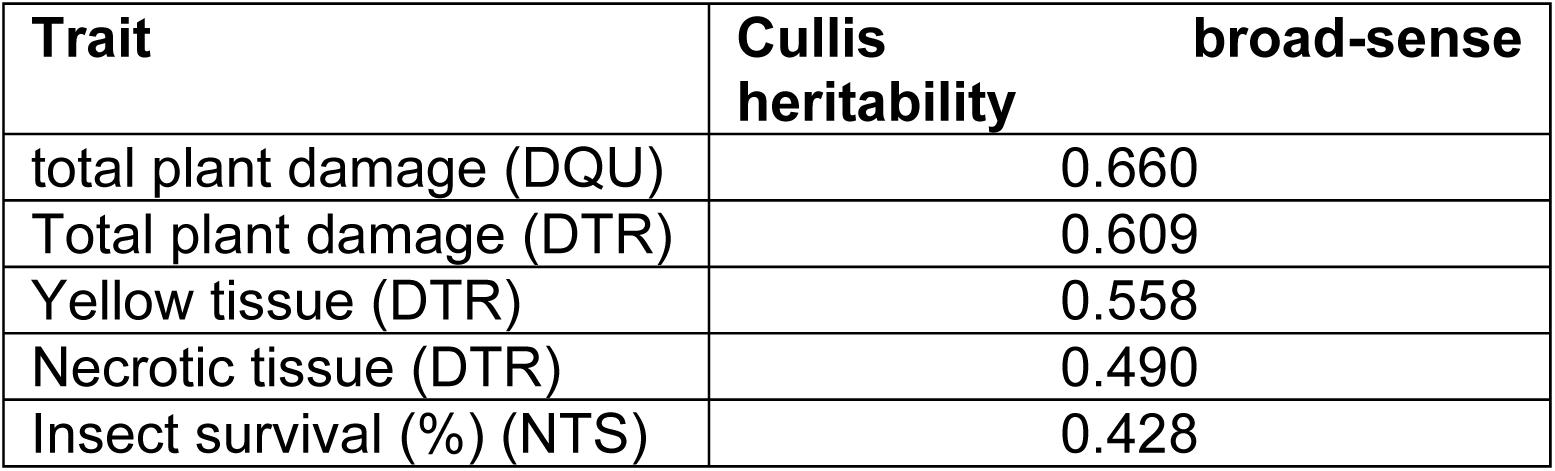
Cullis broad-sense heritability estimates for phenotypic traits in the interspecific *Urochloa* population.

### GWAS Analysis

Principal component analysis (PCA) revealed a clear population structure, with the first two principal components explaining 77.92 % of the total genetic variance (Fig. 5). Individuals clustered according to their biparental family, supporting the presence of stratification consistent with the controlled cross design. To control this structure in the GWAS models, the first three principal components, which cumulatively explained 77.92 % of the genetic variance, were included as covariates in FarmCPU and BLINK models.

**Figure 5.**
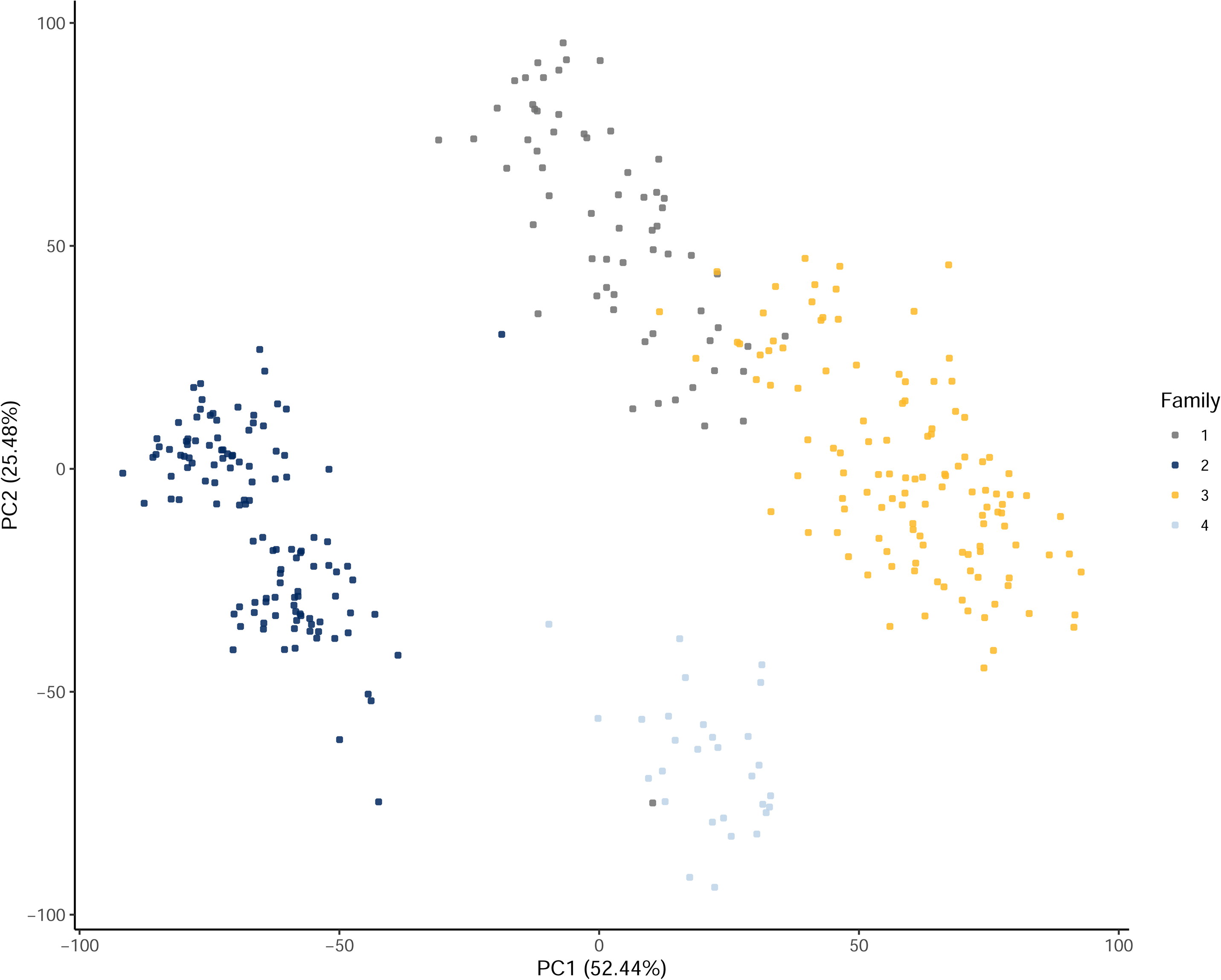
Principal component analysis (PCA) plot from the first two principal components.

Linkage disequilibrium (LD) analysis for the entire F₁ population revealed that LD decayed to r²=0.2 at approximately 2,000 bp. However, substantial variation in LD decay patterns was observed among the four biparental families. While families 1–3 showed similar LD decay distances of ∼1,000 bp, family 4 exhibited extended LD persistence up to ∼3,000 bp. Across chromosomes, LD decay distances (r² < 0.2) ranged from 1 kb to ∼88 kb, with most chromosomes exhibiting decay within 2–10 kb (Fig. 6). Given the heterogeneous LD patterns among families, we opted to use three window sizes (±2 kb, ±5 kb, and ±10 kb) around each significant marker–trait association (MTA) for candidate gene identification.

**Figure 6.**
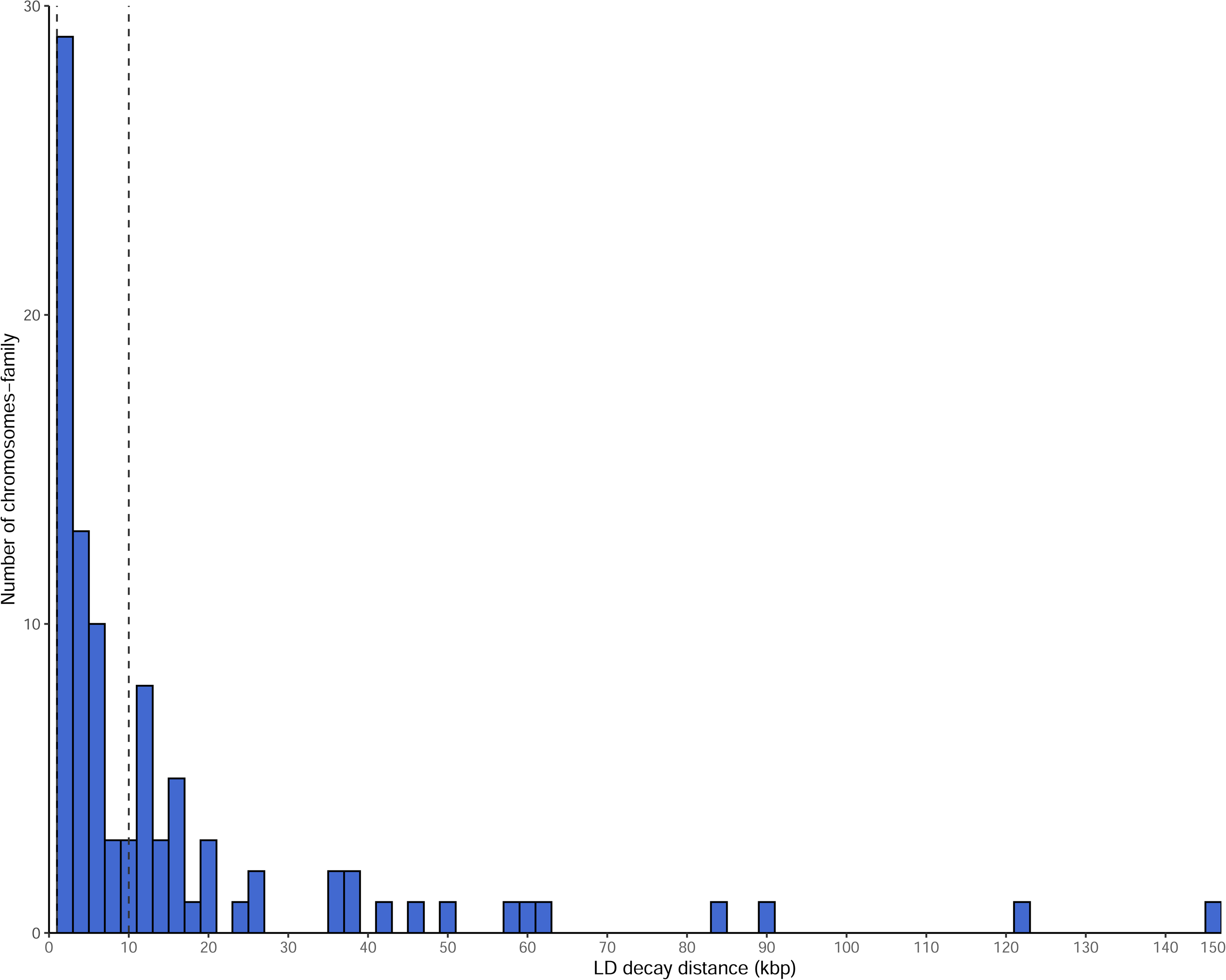
Distribution of LD decay distances (r² = 0.2) across chromosomes in the F₁ mapping population. Vertical dashed lines indicate the 1 kb and 10 kb reference thresholds used to guide QTL window size selection.

Genome-wide association analysis did not identify significant associations with insect survival (NS) but revealed 46 significant marker–trait associations (MTAs) for plant damage phenotypes (Necrotic tissue or total plant damage), all exceeding the threshold of -log_10_(P-value) > 6, using both FarmCPU and BLINK models in GAPIT (Fig. 7; Suppl. Table S2). Q-Q plots demonstrated good model performance with minimal deviation from the expected null distribution and controlled genomic inflation (λ ≈ 1.0), indicating that population structure and kinship were adequately controlled by the inclusion of principal components and kinship matrix as covariates (Fig. 7).

**Figure 7.**
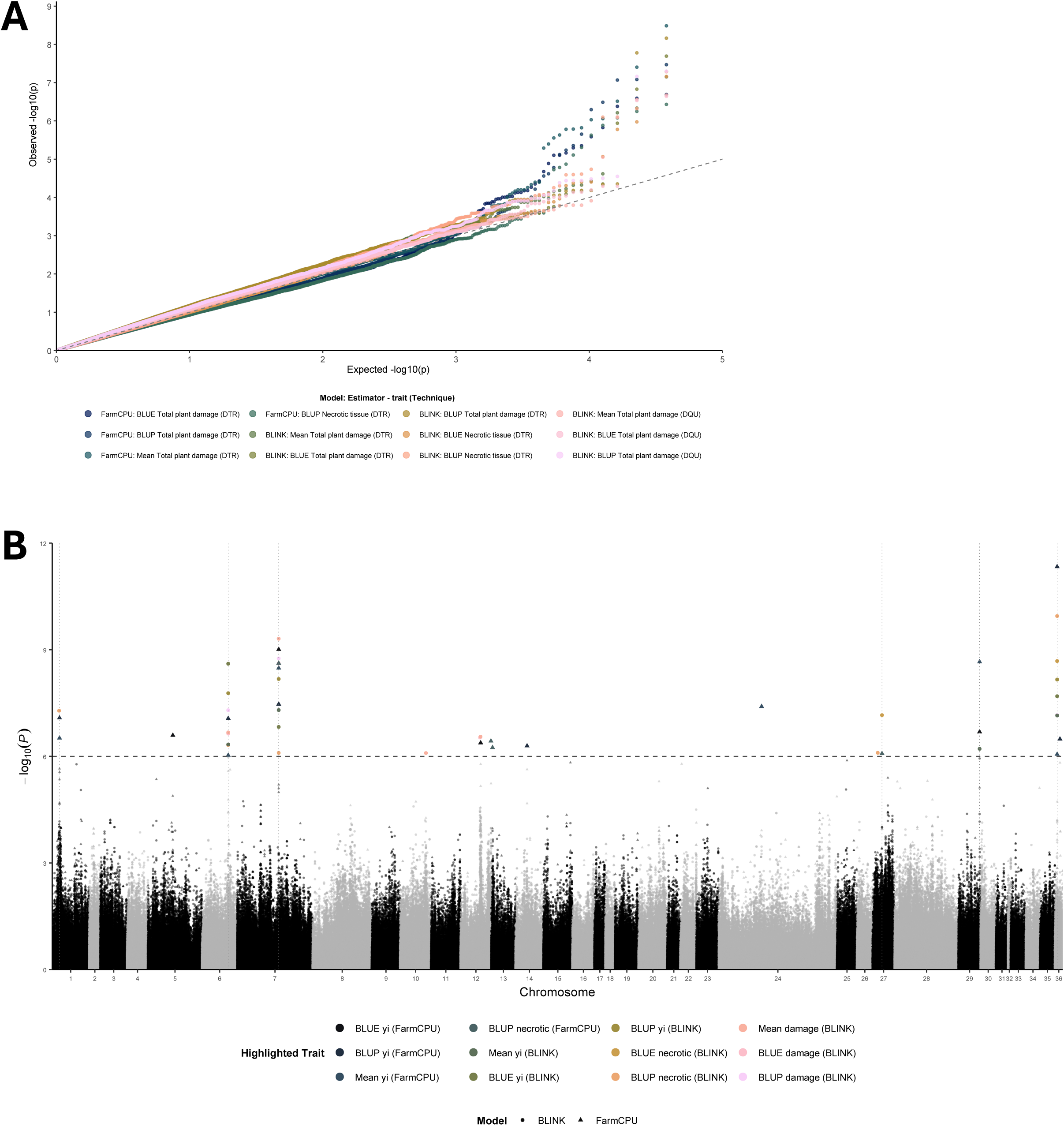
Manhattan plot and QQplot of SNP associations with plant damage traits using BLINK and FarmCPU models. The x-axis represents the genomic position of the markers, and the y-axis shows the –log₁₀(p) values for each SNP. The dashed horizontal line indicates the genome-wide significance threshold (–log₁₀(P-value) = 6). Dotted vertical lines mark chromosomes harboring clusters of significant associations. Different colors and symbols represent trait-model combinations: DTR (damage tolerance rating) and DQU (damage qualification units) assessed using both BLINK and FarmCPU algorithms.

Eighteen consistent and robust QTLs were detected for total plant damage assessed through the DQU methodology, with additional significant associations identified for necrotic tissue damage using the DTR approach (Table 3). Among them, six high-confidence QTLs were located on chromosomes 1, 6, 7, 27, 29, and 36 and consistently detected across both statistical models, multiple phenotypic estimators, and both plant damage quantification methodologies, with overlapping significant MTAs (Table 3). These robust QTLs likely represent genetic variants with stable and reproducible effects on tolerance to *Aeneolamia varia* nymph feeding damage, suggesting multiple independent genetic mechanisms underlying this complex resistance trait. Among these, a locus on chromosome 7 (position 28,519,388) exhibited the largest negative effect size (β = –0.0532) and explained the highest proportion of phenotypic variance (PVE = 21.5%) for spittlebug damage tolerance. However, this locus was associated with a rare favorable allele (MAF = 0.0738), suggesting limited frequency in the studied population. In contrast, a SNP on chromosome 1 (position 11,524,270) displayed a moderate effect (β = 0.0235) coupled with a high allele frequency (MAF = 0.483), making it particularly attractive to try for marker assisted selection in the breeding population. Additional recurrent loci on chromosomes 27, 29, and 36 demonstrated moderate to small effect sizes (β = 0.00903 to 0.0272) with intermediate allele frequencies (MAF = 0.254-0.388), suggesting these variants could contribute additively to spittlebug tolerance.

**Table 3.** Identified QTL for plant damage traits in the interspecific *Urochloa* population.

| Name | Chromosome | Start | End | Trait |
| --- | --- | --- | --- | --- |
| QTL1 | 1 | 11334775 | 11354775 | Necrotic tissue (DTR) |
| QTL2 | 1 | 11514270 | 11534270 | Total plant damage (DQU) |
| QTL3 | 5 | 63376517 | 63396517 | Total plant damage (DQU) |
| QTL4 | 6 | 64514508 | 64534508 | Total plant damage (DQU);<br>Total plant damage (DTR);<br>Necrotic tissue (DTR) |
| QTL5 | 7 | 28509388 | 28529388 | Total plant damage (DQU);<br>Total plant damage (DTR);<br>Necrotic tissue (DTR) |
| QTL6 | 10 | 58791952 | 58811952 | Total plant damage (DTR) |
| QTL7 | 12 | 54702901 | 54722901 | Total plant damage (DTR) |
| QTL8 | 12 | 56104516 | 56124516 | Total plant damage (DTR) |
| QTL9 | 12 | 56374467 | 56394467 | Total plant damage (DQU) |
| QTL10 | 12 | 82648095 | 82668095 | Necrotic tissue (DTR) |
| QTL11 | 13 | 3083711 | 3103711 | Necrotic tissue (DTR) |
| QTL12 | 14 | 23526073 | 23546073 | Total plant damage (DQU) |
| QTL13 | 24 | 52838791 | 52858791 | Total plant damage (DQU) |
| QTL14 | 27 | 13940992 | 13960992 | Necrotic tissue (DTR) |
| QTL15 | 27 | 31335868 | 31355868 | Necrotic tissue (DTR) |
| QTL16 | 29 | 57321694 | 57341694 | Total plant damage (DQU) |
| QTL17 | 36 | 15980330 | 16000330 | Total plant damage (DQU);<br>Total plant damage (DTR);<br>Necrotic tissue (DTR) |
| QTL18 | 36 | 37621131 | 37641131 | Total plant damage (DQU) |

The functional relevance of the identified QTLs was assessed through positional and protein-based annotation. A total of 34 candidate genes were identified within the ±2 kb, ±5 kb, and ±10 kb windows surrounding significant MTAs based on the *Urochloa decumbens* genome annotation (Suppl. Table S3). Initial BLASTp searches against the UniProt database detected 2,101 proteins matches, but after applying stringent filtering criteria (≥85% sequence identity and Protein Existence levels 1 or 2 in UniProt), 30 high-confidence candidate proteins were retained as putative functional genes associated with spittlebug tolerance in the QTL in chromosomes 1, 5, 10, 12, 13, 14 and 27 (Table 4). Among the six high-confidence MTAs identified across both GWAS models, only the SNP on chromosome 1 (position 11,524,270) was within a high-confidence annotated protein, while the remaining five robust MTAs on chromosomes 6,7,27,29 and 36 were located in genomic regions lacking well-annotated protein-coding sequences.

**Table 4.** Significant marker–trait associations (MTAs) for spittlebug damage tolerance identified through GWAS analysis. The table includes GAPIT model parameters, genomic positionsand functional annotated candidate genes with UniProt functional information.

| QTL | *Urochloa* gene name | Start | End | Protein | Main function | Related function with resistance | References |
| --- | --- | --- | --- | --- | --- | --- | --- |
| 1 | LOCUS_993 | 11332693 | 11334978 | Protein ELC-like | Signaling and regulation | Found in the ESCRT complex: cytokinesis, regulation of vesicular trafficking process, cellular autophagy, nuclear envelope reformation, membrane repair, ABA regulation | Spitzer et al., 2006; Isono, 2021; Yu et al., 2020 |
| 1 | LOCUS_994 | 11335110 | 11338139 | Uncharacterized protein | - | - | - |
| 1 | LOCUS_997 | 11345300 | 11348404 | Rac-like GTP-binding protein 5 | Signaling and regulation | Key regulator of immunity responses, cell death and production of ROS | Kawano et al., 2014; Kawasaki et al., 1999 |
| 1 | LOCUS_997 | 11345300 | 11348404 | Rac-like GTP-binding protein 5 |  |  |  |
| 2 | LOCUS_1012 | 11512420 | 11514969 | Glutathione peroxidase | Antioxidant and metabolic enzyme | Eliminate ROS to prevent cellular damage, protect against cell death, ABA signaling pathway, senescence delaying and reinforcement of physical barriers | Santos et al., 2025; Zechmann, 2020 |
| 2 | LOCUS_1012 | 11512420 | 11514969 |  |  |  |  |
| 2 | LOCUS_1012 | 11512420 | 11514969 |  |  |  |  |
| 2 | LOCUS_1013 | 11530189 | 11532991 | Phosphoribosylanthranilate transferase | Antioxidant and metabolic enzyme | Related to shikimate pathway: production of aromatic amino acids such as Tryptophan, production of secondary metabolites involved in plant defense | Lambricht et al., 2013; Scully et al., 2023 |
| 3 | LOCUS_100248 | 63384000 | 63386212 | RING/U-box superfamily protein | Signaling and regulation | Found in UPS: targeting proteins for ubiquitination, regulation of cell-cycle and senescence, regulation of ABA-dependent stomatal responses | Karthik et al., 2025; González-Lamothé et al., 2006 |
| 6 | LOCUS_6034 | 58792240 | 58794245 | Cell wall hydroxyproline-rich glycoprotein | Cell wall structure and barrier | Contribute to the cell wall strengthening and maintain the structure, inhibit the pathogenic cell wall degrading enzymes | Deepak et al., 2010; Yuan-Yuan et al., 2018 |
| 7 | LOCUS_12866 | 54706751 | 54714980 | Oxidoreductase, 2OG-Fe oxygenase family protein | Antioxidant and metabolic enzyme | Involved in oxidation reactions, biosynthesis of phytohormones, jasmonic acid signaling pathway | Wang et al., 2021; Zheng et al., 2022 |
| 7 | LOCUS_12866 | 54706751 | 54714980 | Fe2OG dioxygenase domain-containing protein |  |  |  |
| 8 | LOCUS_12969 | 56112494 | 56115313 | Transmembrane protein | - | - | - |
| 8 | LOCUS_12970 | 56120961 | 56124563 | Anther-specific proline-rich protein APG | Other | Cell wall modification and pollen development. Found in pathogen attack | Roberts et al., 1993; Lee and Cho, 2003 |
| 9 | LOCUS_12990 | 56390023 | 56394977 | F-box protein FBL2 | Signaling and regulation | Found in UPS: targeting proteins for ubiquitination, regulation of cell-cycle and senescence, phytohormones pathway including jasmonic acid | Karthik et al., 2025; Abd-Hamid et al., 2020 |
| 10 | LOCUS_15310 | 82654510 | 82657464 | Very-long-chain aldehyde decarbonylase | Cell wall | Biosynthesis of cuticular waxes | Pascal et al., |
| 10 | LOCUS_<br>15310 | 8265<br>4510 | 8265<br>7464 | aldehyde oxygenase<br>(deformylating) | structure<br>and<br>barrier |  | 2019;<br>Aragón<br>et<br>al.,<br>2021 |
| 11 | LOCUS_<br>15765 | 3097<br>586 | 3101<br>544 | Type I inositol-1,4,5-trisphosphate 5-phosphatase | Signal<br>ing<br>and<br>regula<br>tion | Modulate<br>intracellular calcium<br>levels for signaling<br>defense<br>mechanisms such<br>as movement of<br>stomatal guard<br>cells; involved on<br>systemic acquired<br>resistance | Hung<br>et al.,<br>2014;<br>Perera<br>et al.,<br>2008 |
| 12 | LOCUS_<br>22598 | 2353<br>6761 | 2354<br>0288 | Plant intracellular Ras-group-<br>related LRR protein 4 | Signal<br>ing<br>and<br>regula<br>tion | Signal transduction<br>and pathogen<br>recognition, cell<br>death regulation | Forsth<br>oefel<br>et al.,<br>2005;<br>Lanno<br>o et<br>al.,<br>2014 |
| 14 | LOCUS_<br>68004 | 1394<br>2135 | 1394<br>9209 | Mitosis protein dim1 | Other | Mitotic progression<br>and chromosome<br>segregation, pre-mRNA splicing | Simeo<br>ni and<br>Divita,<br>2007;<br>Berry<br>et al.,<br>1997 |
| 14 | LOCUS_<br>68004 | 1394<br>2135 | 1394<br>9209 | DIM1 |  |  |  |

## Discussion

For over two decades, the interspecific *Urochloa* breeding program at the International Center for Tropical Agriculture (CIAT) has prioritized the evaluation of antibiosis and tolerance as core strategies to breed for spittlebug resistance, achieving substantial genetic gains (Hernández et al., 2022; Worthington & Miles, 2015). Dissecting the genetic architecture underlying these resistance mechanisms in each breeding cycle is, therefore, essential to sustain and enhance these gains, especially considering the extensive genetic polymorphism and environmental plasticity exhibited by spittlebug populations under current and future climate scenarios (Hernández et al., 2021). Our GWAS provide molecular evidence supporting the importance of evaluating both resistance categories independently. The weak phenotypic correlation between insect survival (NTS) and plant damage traits, together with the absence of significant associations for insect survival (antibiosis), contrasted with the identification of 18 significant MTAs for plant damage traits (tolerance), suggests these two mechanisms are controlled by distinct and independent genetic networks. These findings align with the principle that plant resistance to herbivorous insects is often polygenic and continuously shaped by ongoing coevolution with genetically diverse pest populations (Demirjian et al., 2023). Thus, integrating both antibiosis and tolerance mechanisms can contribute to long-term, durable and broad-spectrum resistance in *Urochloa* breeding populations (Priyadarshan, 2019).

The identification of tolerance-specific QTLs provides a foundation for marker-assisted selection approaches that can complement traditional phenotypic evaluation methods, potentially accelerating the development of resilient forage varieties capable of withstanding diverse spittlebug pressures across varying environmental conditions. Insect survival (NTS) exhibited moderate heritability (H² = 0.42), yet no significant SNP associations were identified in our GWAS, highlighting the challenges of detecting genetic signals for traits derived from discrete measurements. In this study, the limited genetic signal observed for NTS likely reflects the statistical complications arising from its derivation from insect counts, which were characterized by non-normal distributions and bounded outcomes constrained to discrete intervals (e.g., 0%, 16.7%, 33.3%). As described by (Reid & Acker, 2022), such distribution reduces statistical power and limits the capacity to fully capture the underlying genetic variance contributing to the trait, consequently, yielding lower estimates of broad-sense heritability. Similar challenges have been reported by (Ferreira et al., 2019), who identified three QTL explaining less than 6.5 % of phenotypic variance in *Notozulia entreriana* nymph survival (H² = 0.37) in a *U. decumbens* intraspecific population with an allele dosage-based linkage mapping approach. The authors attributed these modest effects to the limited resistance of both parentals, insufficient sequencing depth, and methodological constraints intrinsic to reduced-representation genotyping, the last two factors may also have contributed to the lack of NTS-associated SNPs in our study.

The use of single-end reduced-representation sequencing (RAD-seq), while cost-effective for large populations, may have constrained marker density and genotyping resolution, particularly given the segmental allotetraploid nature of *U. decumbens* and the uncharacterized meiotic behavior of the hybrids (Njuguna et al., 2023; Ryan et al., 2025; Worthington et al., 2016). For this reason, standard diploid genotype calling approaches may not adequately capture the true allelic relationships and segregation patterns underlying antibiosis traits, further complicating the detection of genetic associations.

On the other hand, plant damage traits exhibited substantially higher heritability estimates (up to H² = 0.66 for total damage assessed via DQU), supporting the reliability of image-based phenotyping for capturing phenotypic variability within the hybrid *Urochloa* population. The integration of two digital analysis methods, DQU (unsupervised clustering) and DTR (supervised clustering), allowed the quantification of 4,816 images with high precision and speed, without requiring expert evaluation, processing 100 segmented JPEG images in just 4 minutes and 34 seconds (DTR) and 3 minutes and 47 seconds (DQU) on a single processor core. Additionally, the combined use of DTR and DQU methodologies for measuring total plant damage and necrotic areas strengthened the identification of MTAs by providing complementary phenotypic perspectives on tolerance mechanisms. This dual methodology approach enhanced the reliability of our results, as evidenced by the consistency of significant SNP associations detected across both statistical models (FarmCPU and BLINK) and phenotypic methods. This efficiency and scalability are particularly advantageous for GWAS, where precise and continuous phenotypes enhance statistical power to detect MTAs by improving phenotypic resolution across environments and experimental trails in our case, outperforming traditional categorical visual scoring scales in the dissection of complex traits such as tolerance (Araus et al., 2018; McDonald et al., 2022; Xiao et al., 2022).

Overall, and in accordance with previous findings in *Urochloa* for spittlebug resistance (Ferreira et al., 2019) and forage yield-related traits (Vidotti, et al., 2019), the loci associated with tolerance explained moderate proportions of phenotypic variance (PVE < 22%). Among the robust set of six markers, three SNPs on chromosomes 6, 7, and 36 stood out due to their recurrence across multiple models, estimators, and plant damage traits. The SNP on chromosome 7 (position 28,519,388) exhibited the highest phenotypic variance explained (PVE = 21.5%), the largest effect size (β = −0.0532), while being consistently detected across all trait quantifications (DQU, DTR, and necrotic tissue). However, its low minor allele frequency (MAF = 0.07) indicates that the favorable allele is rare within the current breeding population. This potentially limits its applicability by itself in marker-assisted selection, particularly for populations from other breeding cycles where the allele may be rare or absent. In contrast, SNPs on chromosome 6 (position 64,524,508) and chromosome 36 (position 15,990,330) showed moderate phenotypic variance explained and effect (PVE ≈ 5.4%) but higher allele frequencies (MAF = 0.34 and 0.25, respectively), making them more immediately accessible for breeding applications. These moderate-effect loci, while individually contributing smaller phenotypic improvements, offer greater potential for practical implementation due to their higher frequency of favorable alleles and could serve as foundational components in polygenic selection strategies or allele pyramiding approaches.

To harness the full potential of these identified markers, their rigorous validation in diverse genetic contexts is essential for practical application. Thus, future work should focus on the development and validation of Kompetitive Allele-Specific PCR (KASP) markers for the most promising MTAs, as this technology offers a cost-effective and high-throughput method for genotyping diverse breeding populations (Tian et al., 2025). This approach has proven to be successful for resistance to hemipterans in other crops, as demonstrated in cassava for whitefly (*Aleurotrachelus socialis*) resistance by (Bohorquez-Chaux et al., 2025), where consistent GWAS peaks detected across multiple statistical approaches for the same trait, lead to the development of three high-association SNP markers, with one successfully validated for operational marker-assisted selection in the CIAT’s cassava breeding program.

The 18 MTAs identified in this study represent the first set of markers available for spittlebug tolerance in *Urochloa* grasses, providing a foundation for implementing genomic-assisted breeding strategies. Given the moderate effect sizes and varying allele frequencies of these markers, a targeted approach focusing on the most robust associations (particularly those on chromosomes 1, 6, 7, 27, 29 and 36) would be most practical for marker development and validation, e.g. using competitive amplification such as KASP. These validated markers could then be integrated into routine breeding protocols to accelerate the selection of spittlebug-tolerant genotypes and facilitate the pyramiding of favorable alleles across different genetic backgrounds (Worthington et al., 2019).

Our findings support the complex and polygenic nature of spittlebug tolerance in *Urochloa*. As described by (Peterson et al., 2017), insect tolerance is typically governed by multiple loci, each contributing a modest, individual effect through poorly characterized mechanisms, that are often influenced by the intensity of herbivore pressure, and involve a limited number of transcripts functionally linked to resistance at the molecular level. In hemipterans’ resistance systems, mechanisms related to both constitutive and induced resistance have been documented, including compensation of photosynthetic activity, up-regulation of peroxidases and oxidative enzymes and elevated of phytohormones level (Koch et al., 2016).

Candidate proteins functionally related to these established resistance mechanisms were identified in our QTLs, providing valuable insights into their putative roles in the tolerance to *A. varia’s “*feeding damage” in *Urochloa.* These candidates represent a diverse network of protein families spanning multiple functional categories: signaling and regulatory proteins involved in stress perception and response coordination, antioxidant and metabolic enzymes responsible for cellular protection and metabolic adjustment and structural proteins contributing to cell wall reinforcement and physical barriers (Table 4). This functional diversity reflects the multifaceted nature of plant responses to spittlebug herbivory and suggests that tolerance mechanisms in *Urochloa* involve coordinated activation of multiple defense pathways rather than reliance on single resistance genes.

Among the signaling and regulatory proteins, several candidates are likely involved in modulating immune responses, hormone signaling pathways, and stress-induced cell death regulation. More specifically, these proteins are implicated in modulating defense signaling cascades and programmed cell death, as well as coordinating hormone responses, particularly those involving abscisic acid (ABA) and Jasmonic acid (JA)-key phytohormones in the response to herbivory (Abd-Hamid et al., 2020; Forsthoefel et al., 2005; González-Lamothe et al., 2006; Karthik et al., 2025; Kawano et al., 2014; Kawasaki et al., 1999; Lannoo & Van Damme, 2014; Perera et al., 2008; Spitzer et al., 2006; Yu et al., 2020). These proteins often act as intracellular mediators that perceive or transduce signals following herbivory, contributing to plant resistance responses in different stages of pest attack. The identification of hormone signaling components is particularly relevant, as JA-mediated responses are central to induced defenses against chewing insects, while ABA pathways regulate stress tolerance and resource allocation during herbivore pressure (Ballaré, 2011; Wang & Wu, 2013).

The antioxidant and metabolic enzymes group includes candidates with established roles in mitigating oxidative stress, maintaining cellular homeostasis, and supporting the biosynthesis of secondary metabolites. These proteins contribute to detoxifying reactive oxygen species (ROS) generated during herbivore attack, synthesizing defensive phytohormones, and modulating calcium signaling and systemic acquired resistance pathways, which are frequently activated during insect herbivory (do Carmo Santos et al., 2025; Lambrecht & Downs, 2013; Scully et al., 2023; Y. Wang et al., 2021; Zechmann, 2020; Zheng et al., 2022). The presence of oxidative stress response proteins is particularly significant, as spittlebug feeding can disrupt normal cellular processes and generate harmful ROS that must be neutralized to prevent cellular damage and maintain photosynthetic efficiency (Pacheco-Coeto et al., 2019).

In parallel, structural and barrier-related proteins were identified that are likely involved in reinforcing the plant physical defenses. These include enzymes and proteins associated with cell wall fortification, cuticle development, and extracellular matrix stability. By enhancing cell wall strengthening and promoting deposition of protective surface layers, these components serve as a primary line of defense, potentially limiting spittlebug penetration and reducing feeding success(Aragón et al., 2021; Deepak et al., 2010; Pascal et al., 2019; Yuan-Yuan et al., 2018). The identification of cell wall modification enzymes is particularly relevant for spittlebug tolerance, as these insects must penetrate plant tissues to access vascular elements for feeding (Begnami et al., 2025).

Some protein families identified in our study overlap with those reported by (Ferreira et al., 2019) including LRR-containing proteins and F-box domain proteins, both involved in defense signaling and protein degradation pathways. These shared findings reinforce their potential role in both antibiosis and tolerance mechanisms against spittlebugs. However, they also identified WRKY transcription factors, NB-ARC proteins, and pathogenesis-related proteins that were not detected in our analysis, likely reflecting differences in mapping approaches, population genetics, target pest species or the specific resistance mechanisms being evaluated. Conversely, our study identified additional candidates related to oxidative stress mitigation, calcium signaling, and cuticle biosynthesis, suggesting complementary physiological mechanisms specifically involved in *A. varia* nymph tolerance. A few candidate proteins did not show a direct relationship to established plant defense mechanisms. These included proteins involved in general cellular functions such as membrane transport, pollen development, and cell division. For example, one candidate associated with pollen-specific expression and cell wall modification has previously been reported in unrelated plant–pathogen interactions, suggesting a possible role in defense-related signaling (Lee & Cho, 2003; Roberts et al., 1993). Another candidate involved in mitotic progression and pre-mRNA splicing may reflect either baseline cellular maintenance requirements during stress rather than a targeted response to herbivory (Berry & Gould, 1997; Simeoni & Divita, 2007). While their primary functions are not obviously defense-related, these proteins may participate to indirect roles in broader physiological adjustments that support plant survival under herbivore pressure, or they may simply be in linkage disequilibrium with unmapped causal loci.

Additionally, while not all significant SNPs mapped directly to annotated genes, this outcome is expected given the use of reduced representation sequencing (RAD-seq), which captures only a fraction of the genome. In genome-wide association studies, especially those based on RAD-seq, significant SNPs often represent markers in linkage disequilibrium with actual causal variants rather than being causative themselves (Pudjihartono et al., 2022). This is especially relevant in complex polyploid genomes like *Urochloa*, where regulatory elements, non-coding RNAs, or structural variants in intergenic regions may contribute to trait variation. Therefore, these SNPs should be interpreted as valuable markers linked to functional loci that can guide fine-mapping efforts and serve as informative proxies for underlying resistance mechanisms in breeding applications.

The candidate genes identified in this study provide a foundation for several important future research directions that could enhance our understanding of spittlebug tolerance mechanisms and their practical application in the forages breeding program. From a breeding perspective, the functional diversity identified across signaling, metabolic, and structural protein categories suggests that pyramiding alleles from different functional pathways could enhance resistance durability and breadth, potentially providing more robust protection against evolving spittlebug populations and varying environmental pressures (Pilet-Nayel et al., 2017). This multi-pathway approach has shown success in other crop-pest systems, where combining different resistance mechanisms reduces the likelihood of pest adaptation (Poland et al., 2009).

For functional validation priorities, candidates involved in hormone signaling (particularly JA and ABA pathways) and oxidative stress responses should be prioritized, as these represent well-characterized defense mechanisms that can be readily tested through gene expression studies, metabolite profiling, and functional complementation assays. Integration of additional omics approaches, including proteomics to validate protein abundance changes and metabolomics to characterize defensive secondary metabolite profiles, could provide comprehensive validation of these candidate pathways and reveal additional biomarkers for resistance. Furthermore, the identification of SNPs in intergenic regions highlights the potential importance of regulatory variants and non-coding RNAs in controlling tolerance responses, suggesting that future fine-mapping efforts using higher-density markers or whole-genome sequencing could resolve the molecular basis of these associations and identify cis-regulatory elements that modulate candidate gene expression. Such approaches, combined with emerging pan-genome resources, could capture structural variants and regulatory elements missed by current reference-based approaches, providing a more complete picture of the genetic architecture underlying spittlebug tolerance in *Urochloa*

The MTAs identified in this study constitute a valuable contribution for inclusion in the strategic design of a mid-density SNP panel currently under development for genomic selection in the forages breeding program. Rather than relying solely on genome-wide marker distribution, incorporating the 18 spittlebug tolerance-associated SNPs, particularly the six high-confidence markers on chromosomes 1, 6, 7, 27, 29, and 36, into the panel design would ensure that key resistance loci are directly captured rather than relying on linkage disequilibrium with flanking markers. Additionally, incorporating SNPs from the broader QTL regions (±10 kb windows) around each MTA would provide redundancy and capture potential regulatory variants that may contribute to the observed associations. This strategy of combining trait-specific markers with genome-wide coverage has proven effective in other crop genomic selection programs, improving prediction accuracy for target traits while maintaining broad genetic coverage (Anilkumar et al., 2023; Verma et al., 2024).

The integration of these spittlebug tolerance markers into the mid-density panel would enable simultaneous selection for multiple agronomic traits while ensuring that resistance mechanisms are not inadvertently selected against during genomic selection for yield or quality traits. Furthermore, including these validated markers would provide benchmarks for calibrating genomic prediction models and enable the development of weighted selection indices that appropriately balance spittlebug tolerance with other breeding objectives in the *Urochloa* improvement program (Strandén & Jenko, 2024).

Ultimately, these molecular tools and genetic insights will contribute to developing more resilient *Urochloa* cultivars capable of withstanding diverse spittlebug pressures under changing climatic conditions, supporting sustainable livestock production systems across tropical and subtropical regions worldwide.

## Data availability

The digital images used for plant damage quantification are available in the Harvard Dataverse repository at the following identifier: https://dataverse.harvard.edu/dataset.xhtml?persistentId=doi:10.7910/DVN/EGUVHA.

## Acknowledgements

The authors would like to acknowledge the members of the De Vega’s group at the Earlham Institute for their valuable contributions to the development of bioinformatics workflows, as well as the Tropical Forages Program at the International Center for Tropical Agriculture (CIAT) for their ongoing support with phenotypic data collection.

## Literature cited

Abd-Hamid, N. A., Ahmad-Fauzi, M. I., Zainal, Z., & Ismail, I. (2020). Diverse and dynamic roles of F-box proteins in plant biology. In Planta (Vol. 251, Issue 3). Springer. 10.1007/s00425-020-03356-8

Alvarenga, R., Auad, M., Moraes, J. C., Barbosa da Silva, S. E., & Santos Rodrigues, B. (2019). Tolerance to nymphs and adults of Mahanarva spectabilis (Hemiptera: Cercopidae) by forage plants in fertilized soils. Pest Management Science, 75, 2242–2250. 10.1002/ps.5361

Anilkumar, C., Muhammed Azharudheen, T. P., Sah, R. P., Sunitha, N. C., Devanna, B. N., Marndi, B. C., & Patra, B. C. (2023). Gene based markers improve precision of genome-wide association studies and accuracy of genomic predictions in rice breeding. Heredity, 130(5), 335–345. 10.1038/s41437-023-00599-5

Aragón, W., Formey, D., Aviles-Baltazar, N. Y., Torres, M., & Serrano, M. (2021). Arabidopsis thaliana Cuticle Composition Contributes to Differential Defense Response to Botrytis cinerea. Frontiers in Plant Science, 12. 10.3389/fpls.2021.738949

Araus, J. L., Kefauver, S. C., Zaman-Allah, M., Olsen, M. S., & Cairns, J. E. (2018). Translating High-Throughput Phenotyping into Genetic Gain. In Trends in Plant Science (Vol. 23, Issue 5, pp. 451–466). Elsevier Ltd. 10.1016/j.tplants.2018.02.001

Ballaré, C. L. (2011). Jasmonate-induced defenses: A tale of intelligence, collaborators and rascals. In Trends in Plant Science (Vol. 16, Issue 5, pp. 249–257). 10.1016/j.tplants.2010.12.001

Barros, R. D. A., Vital, C. E., Júnior, N. R. S., Vargas, M. A. S., Monteiro, L. P., Faustino, V. A., Auad, A. M., Pereira, J. F., De Oliveira, E. E., Ramos, H. J. O., & Oliveira, M. G. A. (2021). Differential defense responses of tropical grasses to mahanarva spectabilis (Hemiptera: Cercopidae) infestation. Anais Da Academia Brasileira de Ciencias, 93(3). 10.1590/0001-3765202120191456

Bateman, A., Martin, M.-J., Orchard, S., Magrane, M., Adesina, A., Ahmad, S., Bowler-Barnett, E. H., Bye-A-Jee, H., Carpentier, D., Denny, P., Fan, J., Garmiri, P., Gonzales, L. J. da C., Hussein, A., Ignatchenko, A., Insana, G., Ishtiaq, R., Joshi, V., Jyothi, D., … Zhang, J. (2025). UniProt: the Universal Protein Knowledgebase in 2025. Nucleic Acids Research, 53(D1), D609–D617. 10.1093/nar/gkae1010

Begnami, I. dos S., Aono, A. H., Graciano, D. da S., Carmello-Guerreiro, S. M., Ferreira, R. C. U., Malagó, W., Matta, F. de P., Gusmão, M. R., de Souza, A. P., & Vigna, B. B. Z. (2025). Elucidating Molecular Responses to Spittlebug Attack in Paspalum regnellii. Plant Molecular Biology Reporter. 10.1007/s11105-024-01487-w

Berry, L. D., & Gould, K. L. (1997). Fission Yeast dim1 Encodes a Functionally Conserved Polypeptide Essential for Mitosis. In The Journal of Cell Biology (Vol. 137, Issue 6).

Bohorquez-Chaux, A., Becerra Lopez-Lavalle, L. A., Barrera-Enriquez, V., Gómez-Jiménez, M. I., Sanchez-Sarria, C. E., Delgado, L. F., Zhang, X., & Gimode, W. (2025). Genetic mapping and validation of QTL for whitefly resistance in cassava (Manihot esculenta Crantz). Theoretical and Applied Genetics, 138(7). 10.1007/s00122-025-04949-1

Browning, B. L., Tian, X., Zhou, Y., & Browning, S. R. (2021). Fast two-stage phasing of large-scale sequence data. American Journal of Human Genetics, 108(10), 1880–1890. 10.1016/j.ajhg.2021.08.005

Browning, B. L., Zhou, Y., & Browning, S. R. (2018). A One-Penny Imputed Genome from Next-Generation Reference Panels. American Journal of Human Genetics, 103(3), 338–348. 10.1016/j.ajhg.2018.07.015

Butler, D., Cullis, B. R., Gilmour, A. R., Gogel, B. J., & Thompson, R. (2018). ASReml-R Reference Manual Version 4 (4.3.3). VSN International Ltd.

Cardona, C., Miles, J. W., Zuñiga, E., & Sotelo, G. (2010). Independence of Resistance in Brachiaria spp. to Nymphs or to Adult Spittlebugs (Hemiptera: Cercopidae): Implications for Breeding for Resistance. Journal of Economic Entomology, 103(5), 1860–1865. doi: 10.1603/ec10004

Cardona, C., & Sotelo, G. (2005). Mecanismos de resistencia a insectos: naturaleza e importancia en la formulación de estrategias de mejoramiento para incorporar resistencia a salivazo en Brachiaria. Pasturas Tropicales, 27(2), 1–11.

Congio, G. F. S., de Almeida, P. C., Barreto, T. R., Tinazo, V. A., da Silva, T. A. C. C., Costa, D. F. A., & Corsi, M. (2020). Spittlebug damage on tropical grass and its impact in pasture-based beef production systems. Scientific Reports, 10(1), 1–12. 10.1038/s41598-020-67490-9

Covarrubias-Pazaran, G. E. (2020). Heritability: meaning and computation.

da Costa Lima Moraes, A., Mollinari, M., Ferreira, R. C. U., Aono, A., de Castro Lara, L. A., Pessoa-Filho, M., Barrios, S. C. L., Garcia, A. A. F., do Valle, C. B., de Souza, A. P., & Vigna, B. B. Z. (2023). Advances in genomic characterization of Urochloa humidicola: exploring polyploid inheritance and apomixis. Theoretical and Applied Genetics, 136(11), 238. 10.1007/s00122-023-04485-w

Deepak, S., Shailasree, S., Kini, R. K., Muck, A., Mithöfer, A., & Shetty, S. H. (2010). Hydroxyproline-rich glycoproteins and plant defence. In Journal of Phytopathology (Vol. 158, Issue 9, pp. 585–593). 10.1111/j.1439-0434.2010.01669.x

Demirjian, C., Vailleau, F., Berthomé, R., & Roux, F. (2023). Genome-wide association studies in plant pathosystems: success or failure? Trends in Plant Science, 28(4), 471–485. 10.1016/j.tplants.2022.11.006

do Carmo Santos, M. L., Silva Santos, A., Pereira Silva de Novais, D., dos Santos Lopes, N., Pirovani, C. P., & Micheli, F. (2025). The family of glutathione peroxidase proteins and their role against biotic stress in plants: a systematic review. In Frontiers in Plant Science (Vol. 16). Frontiers Media SA. 10.3389/fpls.2025.1425880

Espitia-Buitrago, P., Cotes Torres, J. M., Hernández, L. M., Cardoso, J. A., Chidawanyika, F., & Jauregui, R. N. (2025). Enhancing Phenotyping Accuracy for Selection of Urochloa spp. Tolerant Genotypes to Red Spider Mite (Oligonychus trichardti). Grass and Forage Science, 80(3). 10.1111/gfs.70007

Ferreira, R. C. U., Costa Lima Moraes, A. da, Chiari, L., Simeão, R. M., Vigna, B. B. Z., & de Souza, A. P. (2021). An Overview of the Genetics and Genomics of the Urochloa Species Most Commonly Used in Pastures. In Frontiers in Plant Science (Vol. 12). Frontiers Media S.A. 10.3389/fpls.2021.770461

Ferreira, R. C. U., Lara, L. A. de C., Chiari, L., Barrios, S. C. L., do Valle, C. B., Valério, J. R., Torres, F. Z. V., Garcia, A. A. F., & de Souza, A. P. (2019). Genetic Mapping With Allele Dosage Information in Tetraploid Urochloa decumbens (Stapf) R. D. Webster Reveals Insights Into Spittlebug (Notozulia entreriana Berg) Resistance. Frontiers in Plant Science, 10, 92. 10.3389/fpls.2019.00092

Ferreira, T., & Rasband, W. (2012). ImageJ User Guide ImageJ User Guide IJ 1.46r. https://imagej.net/ij/docs/guide/user-guide.pdf

Forsthoefel, N. R., Cutler, K., Port, M. D., Yamamoto, T., & Vernon, D. M. (2005). PIRLs: A novel class of plant intracellular leucine-rich repeat proteins. Plant and Cell Physiology, 46(6), 913–922. 10.1093/pcp/pci097

González-Lamothe, R., Tsitsigiannis, D. I., Ludwig, A. A., Panicot, M., Shirasu, K., & Jones, J. D. G. (2006). The U-box protein CMPG1 is required for efficient activation of defense mechanisms triggered by multiple resistance genes in tobacco and tomato. Plant Cell, 18(4), 1067–1083. 10.1105/tpc.106.040998

Hernández, L., Espitia, P., & Cardoso, J. A. (2022). Digital imaging outperforms traditional scoring methods for spittlebug tolerance in Urochloa humidicola hybrids. Tropical Grasslands-Forrajes Tropicales, 10(3), 271–279. 10.17138/TGFT(10)271-279

Hernández, L. M., Espitia, P., Florian, D., Castiblanco, V., Cardoso, J. A., & Gómez-Jiménez, M. I. (2021). Geographic Distribution of Colombian Spittlebugs (Hemiptera: Cercopidae) via Ecological Niche Modeling: A Prediction for the Main Tropical Forages’ Pest in the Neotropics. Frontiers in Sustainable Food Systems, 5(725774). 10.3389/fsufs.2021.725774

Hernández, L. M., Sandoval, K., Aparicio, J., Ariza, D., Espitia-Buitrago, P., Castiblanco, V., & Jauregui, R. N. (2022, September 14). Estimation of genetic gain for resistance to spittlebugs (Hemiptera: Cercopidae) in the Interspecific Urochloa CIAT Breeding Program using historical data. Presentation Prepared for Tropentag 2022 - Can Agroecological Farming Feed the World? Farmers’ and Academia’s Views.

Huang, M., Liu, X., Zhou, Y., Summers, R. M., & Zhang, Z. (2019). BLINK: A package for the next level of genome-wide association studies with both individuals and markers in the millions. GigaScience, 8(2). 10.1093/gigascience/giy154

Karthik, H. N., Parmar, S., Gawande, N. D., & Sankaranarayanan, S. (2025). Multifaceted Roles of U-Box E3 Ligases in Plant Development. Plant and Cell Physiology. 10.1093/pcp/pcaf059

Kawano, Y., Kaneko-Kawano, T., & Shimamoto, K. (2014). Rho family GTPase-dependent immunity in plants and animals. Frontiers in Plant Science, 5(OCT). 10.3389/fpls.2014.00522

Kawasaki, T., Henmi, K., Ono, E., Hatakeyama, S., & Shimamoto, K. O. (1999). The small GTP-binding protein Rac is a regulator of cell death in plants (Vol. 96). www.pnas.org.

Koch, K. G., Chapman, K., Louis, J., Heng-Moss, T., & Sarath, G. (2016). Plant tolerance: A unique approach to control hemipteran pests. Frontiers in Plant Science, 7: 1363(September). 10.3389/fpls.2016.01363

Lambrecht, J. A., & Downs, D. M. (2013). Anthranilate phosphoribosyl transferase (TrpD) generates phosphoribosylamine for thiamine synthesis from enamines and phosphoribosyl pyrophosphate. ACS Chemical Biology, 8(1), 242–248. 10.1021/cb300364k

Langmead, B., & Salzberg, S. L. (2012). Fast gapped-read alignment with Bowtie 2. Nature Methods, 9(4), 357–359. 10.1038/nmeth.1923

Lannoo, N., & Van Damme, E. J. M. (2014). Lectin domains at the frontiers of plant defense. In Frontiers in Plant Science (Vol. 5, Issue AUG). Frontiers Research Foundation. 10.3389/fpls.2014.00397

Lee, K.-A., & Cho, T.-J. (2003). Characterization of a Salicylic Acid-and Pathogen-induced Lipase-like Gene in Chinese Cabbage. Journal of Biochemistry and Molecular Biology, 36(5), 433–441. http://www.cbs.dtu.dk/

Liu, X., Huang, M., Fan, B., Buckler, E. S., & Zhang, Z. (2016). Iterative Usage of Fixed and Random Effect Models for Powerful and Efficient Genome-Wide Association Studies. PLoS Genetics, 12(2). 10.1371/journal.pgen.1005767

Matias, F. I., Vidotti, M. S., Meireles, K. G. X., Barrios, S. C. L., Do Valle, C. B., Carley, C. A. S., & Fritsche-Neto, R. (2019). Association mapping considering allele dosage: An example of forage traits in an interspecific segmental allotetraploid urochloa spp. panel. Crop Science, 59(5), 2062–2076. 10.2135/cropsci2019.03.0185

Matias, F. I., Xavier Meireles, K. G., Nagamatsu, S. T., Lima Barrios, S. C., Borges do Valle, C., Carazzolle, M. F., Fritsche-Neto, R., & Endelman, J. B. (2019). Expected Genotype Quality and Diploidized Marker Data from Genotyping-by-Sequencing of Urochloa spp. Tetraploids. The Plant Genome, 12(3), 190002. 10.3835/plantgenome2019.01.0002

McDonald, S. C., Buck, J., & Li, Z. (2022). Automated, image-based disease measurement for phenotyping resistance to soybean frogeye leaf spot. Plant Methods, 18(1). 10.1186/s13007-022-00934-7

Nitthaisong, P., Ishigaki, G., Suenaga, K., Muguerza, M., Tanaka, H., & Akashi, R. (2019). Pentaploid apomicts by interspecific hybridization between diploid urochloa ruziziensis and tetraploid apomictic U. Decumbens. Crop Science, 59(4), 1648–1656. 10.2135/cropsci2019.01.0035

Njuguna, J. N., Clark, L. V., Lipka, A. E., Anzoua, K. G., Bagmet, L., Chebukin, P., Dwiyanti, M. S., Dzyubenko, E., Dzyubenko, N., Ghimire, B. K., Jin, X., Johnson, D. A., Kjeldsen, J. B., Nagano, H., de Bem Oliveira, I., Peng, J., Petersen, K. K., Sabitov, A., Seong, E. S., … Sacks, E. J. (2023). Impact of genotype-calling methodologies on genome-wide association and genomic prediction in polyploids. Plant Genome, 16(4). 10.1002/tpg2.20401

Pacheco-Coeto, R., Cárdenas-Torres, L., Hernández-Rosas, F., Valente Hidalgo-Contreras, J., & Aquino-Pérez, G. (2019). Callose and reactive oxygen species expressed in sugar cane leaves by mechanical damage of spittlebugs. Rev. Mex. Cienc. Agríc. Esp. Pub. Num, 22.

Pamidi, L. S., Sekhar, J. C., Yathish, K. R., Karjagi, C. G., Rao, K. S., Suby, S. B., & Rakshit, S. (2025). Characterization of antixenosis and antibiosis resistance to the fall armyworm, Spodoptera frugiperda (J.E. Smith) in maize. Phytoparasitica, 53(1). 10.1007/s12600-024-01228-5

Parsa, S., Sotelo, G., & Cardona, C. (2011). Characterizing herbivore resistance mechanisms: Spittlebugs on Brachiaria spp. as an example. Journal of Visualized Experiments, 52. 10.3791/3047

Pascal, S., Bernard, A., Deslous, P., Gronnier, J., Fournier-Goss, A., Domergue, F., Rowland, O., & Joubès, J. (2019). Arabidopsis CER1-LIKE1 functions in a cuticular very-long-chain alkane-forming complex. Plant Physiology, 179(2), 415–432. 10.1104/pp.18.01075

Patrignani, A., & Ochsner, T. E. (2015). Canopeo: A powerful new tool for measuring fractional green canopy cover. Agronomy Journal, 107(6), 2312–2320. 10.2134/agronj15.0150

Perea, C., De La Hoz, J. F., Cruz, D. F., Lobaton, J. D., Izquierdo, P., Quintero, J. C., Raatz, B., & Duitama, J. (2016). Bioinformatic analysis of genotype by sequencing (GBS) data with NGSEP. BMC Genomics, 17. 10.1186/s12864-016-2827-7

Perera, I. Y., Hung, C. Y., Moore, C. D., Stevenson-Paulik, J., & Boss, W. F. (2008). Transgenic Arabidopsis plants expressing the type 1 inositol 5-phosphatase exhibit increased drought tolerance and altered abscisic acid signaling. Plant Cell, 20(10), 2876–2893. 10.1105/tpc.108.061374

Peterson, R. K. D., Varella, A. C., & Higley, L. G. (2017). Tolerance: The forgotten child of plant resistance. PeerJ, 5: *e3934*. 10.7717/peerj.3934

Pilet-Nayel, M. L., Moury, B., Caffier, V., Montarry, J., Kerlan, M. C., Fournet, S., Durel, C. E., & Delourme, R. (2017). Quantitative resistance to plant pathogens in pyramiding strategies for durable crop protection. In Frontiers in Plant Science (Vol. 8). Frontiers Media S.A. 10.3389/fpls.2017.01838

Poland, J. A., Balint-Kurti, P. J., Wisser, R. J., Pratt, R. C., & Nelson, R. J. (2009). Shades of gray: the world of quantitative disease resistance. In Trends in Plant Science (Vol. 14, Issue 1, pp. 21–29). 10.1016/j.tplants.2008.10.006

Priyadarshan, P. M. (2019). Host Plant Resistance Breeding. In Plant breeding: Classical to Modern (1st ed., pp. 379–412). Springer Nature Singapore. 10.1007/978-981-13-7095-3_18

Pudjihartono, N., Fadason, T., Kempa-Liehr, A. W., & O’Sullivan, J. M. (2022). A Review of Feature Selection Methods for Machine Learning-Based Disease Risk Prediction. Frontiers in Bioinformatics, 2. 10.3389/fbinf.2022.927312

Reid, J. M., & Acker, P. (2022). Properties of phenotypic plasticity in discrete threshold traits. Evolution, 76(2), 190–206. 10.1111/evo.14408

Roberts, M. R., Foster, G. D., Blundell, R. P., Robinson, S. W., Kumar, A., Draper, J., & Scott, R. (1993). Gametophytic and sporophytic expression of an antherspecific Arabidopsis thaliana gene . The Plant Journal, 3(1), 111–120. 10.1046/j.1365-313x.1993.t01-5-00999.x

Ryan, C., Fraser, F., Irish, N., Barker, T., Knitlhoffer, V., Durrant, A., Reynolds, G., Kaithakottil, G., Swarbreck, D., & De Vega, J. J. (2025). A haplotype-resolved chromosome-level genome assembly of *Urochloa decumbens* cv. Basilisk resolves its allopolyploid ancestry and composition. G3: Genes, Genomes, Genetics. 10.1093/g3journal/jkaf005

Scully, T. W., Jiao, W., Mittelstädt, G., & Parker, E. J. (2023). Structure, mechanism and inhibition of anthranilate phosphoribosyltransferase. In Philosophical Transactions of the Royal Society B: Biological Sciences (Vol. 378, Issue 1871). Royal Society Publishing. 10.1098/rstb.2022.0039

Silva, S. E. B., Auad, A. M., Moraes, J. C., Alvarenga, R., Claudino, S. S., & Resende, T. T. (2017). Biological performance and preference of Mahanarva spectabilis (Hemiptera: Cercopidae) for feeding on different forage plants. Journal of Economic Entomology, 110(4), 1877–1885. 10.1093/jee/tox180

Silva, S. E. B., Auad, A. M., Moraes, J. C., Alvarenga, R., Fonseca, M. G., Marques, F. A., Santos, N. C. S., & Nagata, N. (2019). Olfactory response of Mahanarva spectabilis (Hemiptera: Cercopidae) to volatile organic compounds from forage grasses. Scientific Reports, 9(1), 1–6. 10.1038/s41598-019-46693-9

Simeoni, F., & Divita, G. (2007). The Dim protein family: From structure to splicing. In Cellular and Molecular Life Sciences (Vol. 64, Issue 16, pp. 2079–2089). 10.1007/s00018-007-7043-9

Souza, B. H. S. (2025). Host Plant Resistance: Is It Time for a New Model? Neotropical Entomology, 54(1). 10.1007/s13744-025-01300-7

Spitzer, C., Schellmann, S., Sabovljevic, A., Shahriari, M., Keshavaiah, C., Bechtold, N., Herzog, M., Müller, S., Hanisch, F. G., & Hülskamp, M. (2006). The Arabidopsis elch mutant reveals functions of an ESCRT components in cytokinesis. Development, 133(23), 4679–4689. 10.1242/dev.02654

Stout, M. J., Bernaola, L., & Acevedo, F. (2024). Recent history and future trends in host-plant resistance. In Annals of the Entomological Society of America (Vol. 117, Issue 3, pp. 139–149). Entomological Society of America. 10.1093/aesa/saae006

Strandén, I., & Jenko, J. (2024). A computationally feasible multi-trait single-step genomic prediction model with trait-specific marker weights. Genetics Selection Evolution, 56(1). 10.1186/s12711-024-00926-2

Tian, Y., Liu, P., Kong, D., Nie, Y., Xu, H., Han, X., Sang, W., & Li, W. (2025). Genome-wide association analysis and KASP markers development for protein quality traits in winter wheat. BMC Plant Biology, 25(1). 10.1186/s12870-025-06171-z

Urbanek, S. (2011). jpeg: Read and write JPEG images (R package version 0.1-11). https://cran.r-project.org/web/packages/jpeg/jpeg.pdf

VanRaden, P. M. (2008). Efficient methods to compute genomic predictions. Journal of Dairy Science, 91(11), 4414–4423. 10.3168/jds.2007-0980

Verma, S., Gupta, A. R. S. S. H., Yalla, S., Shreya, Patel, P. J., Sharma, R., & Donga, A. (2024). Integrating Marker-Assisted (MAS) and Genomic Selection (GS) for Plant Functional Trait Improvement. In N. Kumar & H. Singh (Eds.), Plant Functional Traits for Improving Productivity (pp. 203–215). Springer Nature Singapore.

Wang, J., & Zhang, Z. (2021). GAPIT Version 3: Boosting Power and Accuracy for Genomic Association and Prediction. *Genomics*, Proteomics and Bioinformatics, 19(4), 629–640. 10.1016/j.gpb.2021.08.005

Wang, L., & Wu, J. (2013). The Essential Role of Jasmonic Acid in Plant-Herbivore Interactions - Using the Wild Tobacco Nicotiana attenuata as a Model. In Journal of Genetics and Genomics (Vol. 40, Issue 12, pp. 597–606). 10.1016/j.jgg.2013.10.001

Wang, Y., Shi, Y., Li, K., Yang, D., Liu, N., Zhang, L., Zhao, L., Zhang, X., Liu, Y., Gao, L., Xia, T., & Wang, P. (2021). Roles of the 2-oxoglutarate-dependent dioxygenase superfamily in the flavonoid pathway: A review of the functional diversity of f3h, fns i, fls, and ldox/ans. In Molecules (Vol. 26, Issue 21). MDPI. 10.3390/molecules26216745

Worthington, M., Ebina, M., Yamanaka, N., Heffelfinger, C., Quintero, C., Zapata, Y. P., Perez, J. G., Selvaraj, M., Ishitani, M., Duitama, J., De la Hoz, J. F., Rao, I., Dellaporta, S., Tohme, J., & Arango, J. (2019). Translocation of a parthenogenesis gene candidate to an alternate carrier chromosome in apomictic Brachiaria humidicola. BMC Genomics, 20(1). 10.1186/s12864-018-5392-4

Worthington, M., Heffelfinger, C., Bernal, D., Quintero, C., Zapata, Y. P., Perez, J. G., De Vega, J., Miles, J., Dellaporta, S., & Tohme, J. (2016). A parthenogenesis gene candidate and evidence for segmental allopolyploidy in apomictic Brachiaria decumbens. In Genetics (Vol. 203). 10.1534/genetics.116.190314

Worthington, M. L., & Miles, J. W. (2015). Reciprocal Full-sib Recurrent Selection and Tools for Accelerating Genetic Gain in Apomictic Brachiaria. In H. Budak & G. Spangenber (Eds.), Molecular Breeding of Forage and Turf (pp. 111–122). Springer International Publishing Switzerland 2015. 10.1007/978-3-319-08714-6_10

Worthington, M., Perez, J. G., Mussurova, S., Silva-Cordoba, A., Castiblanco, V., Cardoso Arango, J. A., Jones, C., Fernandez-Fuentes, N., Skot, L., Dyer, S., Tohme, J., DI Palma, F., Arango, J., Armstead, I., & De Vega, J. J. (2021). A new genome allows the identification of genes associated with natural variation in aluminium tolerance in Brachiaria grasses. Journal of Experimental Botany, 72(2), 302–319. 10.1093/jxb/eraa469

Xiao, Q., Bai, X., Zhang, C., & He, Y. (2022). Advanced high-throughput plant phenotyping techniques for genome-wide association studies: A review. In Journal of Advanced Research (Vol. 35, pp. 215–230). Elsevier B.V. 10.1016/j.jare.2021.05.002

Yu, F., Cao, X., Liu, G., Wang, Q., Xia, R., Zhang, X., & Xie, Q. (2020). ESCRT-I Component VPS23A Is Targeted by E3 Ubiquitin Ligase XBAT35 for Proteasome-Mediated Degradation in Modulating ABA Signaling. Molecular Plant, 13(11), 1556–1569. 10.1016/j.molp.2020.09.008

Yuan-Yuan, L., Chen, X. M., Zhang, Y., Cho, Y. H., Wang, A. R., Yeung, E. C., Zeng, X., Guo, S. X., & Lee, Y. I. (2018). Immunolocalization and changes of hydroxyproline-rich glycoproteins during symbiotic germination of dendrobium officinale. Frontiers in Plant Science, 9. 10.3389/fpls.2018.00552

Zechmann, B. (2020). Subcellular roles of glutathione in mediating plant defense during biotic stress. In Plants (Vol. 9, Issue 9, pp. 1–21). MDPI AG. 10.3390/plants9091067

Zhang, C., Dong, S. S., Xu, J. Y., He, W. M., & Yang, T. L. (2019). PopLDdecay: A fast and effective tool for linkage disequilibrium decay analysis based on variant call format files. Bioinformatics, 35(10), 1786–1788. 10.1093/bioinformatics/bty875

Zheng, H., Wang, Y., Li, X., Huang, W., Miao, H., Li, H., Wang, Q., Sun, B., & Zhang, F. (2022). A novel putative 2-oxoglutarate-dependent dioxygenase gene (BoaAOP-like) regulates aliphatic glucosinolate biosynthesis in Chinese kale. Scientia Horticulturae, 297. 10.1016/j.scienta.2022.110921

Zhiwu Zhang Laboratory. (2025). User Manual for GAPIT: Genomic Association and Prediction Integrated Tool (Version 3). http://zzlab.net/GAPIT

